# RORγt serine 182 tightly regulates T cell heterogeneity to maintain mucosal homeostasis and restrict tissue inflammation

**DOI:** 10.1101/2021.05.14.444071

**Authors:** Shengyun Ma, Shefali Patel, Nicholas Chen, Parth R. Patel, Benjamin S. Cho, Nazia Abbasi, John T. Chang, Wendy Jia Men Huang

## Abstract

Intestine homeostasis is maintained by the delicate balance of Th17 effector cells and Treg cells. Dysregulation of these cell populations contributes to inflammation, tissue damage, and chronic conditions. RORγt is essential for the differentiation of Th17 and a subset of Treg (RORγt^+^ Treg) cells involved in intestinal inflammation. RORγt belongs to the nuclear receptor family of transcription factors with hinge regions that are highly flexible for co-activator/co-repressor interactions. Serine 182 at the hinge region of RORγt is phosphorylated. This study aims to uncover how S182 on RORγt contributes to mucosal homeostasis and diseases. We used CRISRP technology to generate a phosphor-null knock-in mutant mouse line (RORγt^S182A^) to assess its role in intestine physiology. scRNA-seq was performed on WT and RORγt^S182A^ cohoused littermates to evaluate colonic T cell heterogeneity under steady state and colitis settings. Single-cell transcriptomics revealed that RORγt^S182^ maintains colonic T cell heterogeneity under steady state, without interfering T cell development and differentiation. In inflamed tissues, RORγt^S182^ simultaneously restricts IL-1β-mediated Th17 activities and promotes anti-inflammatory cytokine IL-10 production in LT-like Treg cells. Phospho-null RORγt^S182A^ knock-in mice challenged with DSS induced colitis and EAE experienced delayed recovery and exacerbated pathology. The double switch role of RORγt^S182^ is critical in resolving T cell-mediated inflammation and provides a potential therapeutic target to combat autoimmune diseases.

## INTRODUCTION

The *Rorc* locus encodes two isoforms of RORγ, an evolutionarily conserved member of the nuclear receptor transcription factor family. The long isoform (RORγ) is broadly expressed in the liver, kidney, and muscles, and the shorter isoform (RORγt) is uniquely found in cells of the immune system ^1^. In the thymus, RORγt promotes the expression of the anti-apoptotic factor, B-cell lymphoma-extra large (Bcl-xL), to prevent apoptosis of CD4^+^CD8^+^ cells during T cell development ^2, 3^. In the periphery, RORγt is essential for secondary lymphoid tissue organogenesis ^4^ and the differentiation of effector T lymphocyte and ILCs involved in mucosal protection ^5–7^. Humans with loss of function mutations of *RORC* have impaired anti-bacterial and anti-fungal immunity ^8^. More importantly, RORγt is the master transcription factor required for the production of cytokines implicated in protection against microbial infections, as well as autoimmune disease pathogenesis, including interleukin 17 (IL-17) cytokines in pathogenic Th17 cells and IL-10 in RORγt^+^Treg cells ^6, 9–12^. However, the mechanisms by which RORγt controls diverse programs in a cell-type-specific manner are not well understood, thus, limiting our ability to target this pathway to ameliorate autoimmune problems without adversely affecting other homeostatic processes that are also highly dependent on RORγt.

The RORγt protein consists of the DBD, a hinge region, and a LBD. PTMs at the DBD and LBD have been reported to modulate RORγt functions ^13^. Lysine acetylations at the DBD regulate RORγt binding to chromatin DNA ^*14*^ and lysine ubiquitinations at the DBD regulate RORγt protein turnover ^15–17^. Serine phosphorylations at the LBD fine-tune RORγt transcription activities at the *Il17a* locus in Th17 cells ^18^ and contribute to autoimmunity ^19^. However, little is known about the contribution of the highly conserved RORγt hinge region and its associated PTMs in its diverse *in vivo* functions.

Here, we report that RORγt is phosphorylated at serine 182 at the hinge region. Importantly, this residue is dispensable for T cell development in the thymus and T effector cell differentiation in the periphery. scRNA-seq revealed that RORγt^S182^ maintains colonic T cell heterogeneity under steady state. In inflamed tissues, RORγt^S182^ is a critical negative feedback mechanism for restricting IL-1β-mediated Th17 cell effector functions, but also acts to promote anti-inflammatory cytokine IL-10 production in LT-like RORγt^+^ Treg cells. Phosphor-null RORγt^S182A^ knock-in mice experienced delayed recovery and succumbed to more severe diseases after DSS induced colitis and EAE challenges.

## RESULTS

### Serine 182 of RORγt is dispensable for thymic T cell development

To characterize PTMs of RORγt, a 2xFlag-tagged murine RORγt expression construct was transfected into HEK293ft cells. Whole-cell lysates after transfection 48hrs were used for anti-Flag immunoprecipitation and tandem mass-spectrometry analysis. Peptide mapping to RORγt harbored phosphorylation at S182, T197, S489, methylation at R185, and ubiquitination at K495 (Figure 1A). In particular, the serine residue in position 182 of RORγt is evolutionarily conserved between mouse and human (Supplementary Figure 1A). Phosphorylation at this site was the most abundant among all the RORγt PTMs identified (Figure 1B), consistent with previous reports ^18, 25^.

**Figure 1.**
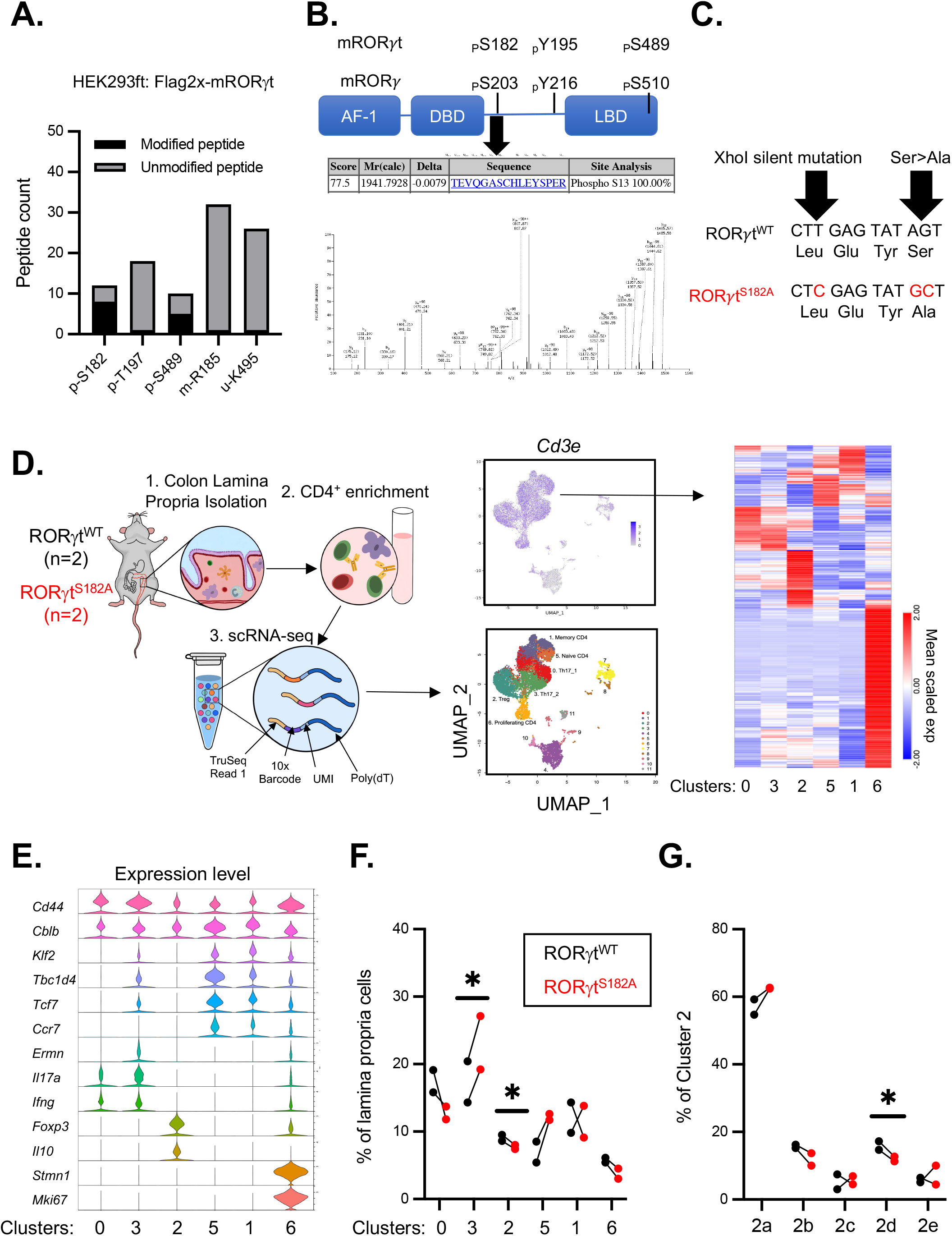
RORγt^S18^-dependent Th17 and Treg populations in steady state colon. A. Proportion of modified and unmodified peptides identified by Tandem MS/MS mapping to murine RORγt from whole cell lysates of 293ft cells transfected with Flag2x-mRORγt expression construct for 48hrs. B. Top: Major phosphorylation sites in relation to the activation function (AF-1), DBD and LBD of RORγt. Bottom: representative mass spectral map of one murine RORγt peptide carrying a phosphorylated S182 residue. C. CRISPR-Cas mediated genomic mutations to generate the RORγt^S182A^ knock-in mice. D. Left: scRNA-seq experiment workflow. Center: UMAP plot of twelve immune cell clusters obtained from D (left) and the average expression of *Cd3e* in each cell (right). Right: Heatmap of average expression of genes with cluster-specific patterns. E. Violin plots of selected genes from each cluster, with gene expression displayed on the y-axis. F. Proportions of colonic CD4^+^ T cells in each cluster from two pairs of WT and RORγt^S182A^ littermates. G. Proportions of Treg subclusters from steady state colons of WT and RORγt^S182A^ mice (n=2). * p-value<0.05 (multiple t-test).

To assess the *in vivo* function of this serine residue, we generated a phospho-null knock-in mice line (RORγt^S182A^) by replacing the serine codon on *Rorc* with that of alanine using CRISPR-Cas9 technology (Figure 1C). Heterozygous crosses yielded WT and homozygous RORγt^S182A^ littermates born in Mendelian ratio with similar growth rates (Supplementary Figure 1B). In RORγt^S182A^ CD4^+^CD8^+^ thymocytes, RORγt and Bcl-xL protein abundance were similar to those observed in WT cells (Supplementary Figure 1C-D). RORγt^S182A^ CD4^+^CD8^+^ thymocytes matured normally into single positive CD4^+^ T helper and CD8^+^ cytotoxic T cells (Supplementary Figure 1E). Mature CD4^+^ T helper and CD8^+^ cytotoxic cell populations in the spleen also appeared normal in the RORγt^S182A^ mice (Supplementary Figure 1F). Together, these results suggest that serine 182 and/or its phosphorylation is not involved in thymic RORγt protein turnover and is dispensable for T cell development *in vivo*.

### scRNA-seq revealed RORγt^S182^ dependent Th17 and Treg programs in the steady state colon

In the intestinal lamina propria, RORγt is required for the differentiation of Th17 and RORγt^+^ Treg cells, and ILC3 ^6, 26, 27^ To our surprise, a similar proportion of RORγt-expressing CD4^+^ T cells and ILC3 (CD3^-^) cells were present in both the small intestine and colon of WT and RORγt^S182A^ mice (Supplementary Figure 1G-H), suggesting that serine 182 on RORγt is dispensable for their generation. Next, we asked whether serine 182 on RORγt regulates T cell heterogeneity and subset specific gene programs. The single-cell RNA transcriptome of 10,000 - 13,000 colonic lamina propria CD4^+^ T cells from two pairs of WT and RORγt^S182A^ mice were profiled according to manufacturer instructions (10x Genomics) and analyzed using Seraut ^28^. Uniform manifold approximation and projection (UMAP) of the 11,725 cells with *Cd4*>0.4 and ~1,100 genes/cell revealed 12 clusters. We performed in-depth analysis on 6 clusters with high co-expression of *Cd4* and *Cd3*, which consists of 8,175 cells (69.7% total) (outlined in Figure 1D).

Population level analysis suggests that WT and RORγt^S182A^ colons had similar proportions of memory T cells (cluster 1, *Cd44*^hi^). In the RORγt^S182A^ colon, there was an increase in the proportion of naïve T cells (cluster 5, *Cd44*^low^), a reduction in the proportion of actively proliferating T cells (cluster 6, *Ki67*^high^) and Treg cells (cluster 2, *Foxp3*^high^ and *Il10*^high^) (Figure 1E-F). Among the five Treg subsets, subsets 2a and 2e expressed genes characteristics of suppressive Treg cells, including *Gzmb* and *Ccr1* (Supplementary Figure 2A-B), as described in the previous report ^29^. Subsets 2c and 3d expressed genes characteristics of LT-like Treg, including *S1pr1*. Subset 2b expressed genes characteristics of non-lymphoid tissues Treg, including *Gata3* and *Pdcd1*. RORγt^S182A^ colon harbored a reduced proportion of LT-like Tregs (cluster 2d) (Figure 1G).

scRNA-seq revealed two main effector T cell clusters (0 and 3) as two closely related subsets of the Th17 lineage with marked expression of *Il17a* and *Ifng* (Figure 1E). Cluster 0 Th17 cells expressed greater *Il18r1, Il1rn, Ccl2, Ccl5, Ccl7, Ccl8, Cxcl1, and Cxcl13*. In contrast, Th17 cells in cluster 3 had higher levels of *Il23r, Ccr6, Il22, Ccl20*, and *Cxcl3* (Supplementary Figure 2C). In WT colon, the two Th17 subsets were found in a balanced 1:1 ratio. This balance was significantly disrupted in the RORγt^S182A^ colon, where cluster 3 Th17 cells dominate (Figure 1F). These scRNA-seq results reveal an unexpected role of RORγt^S182^ in maintaining colonic T cell heterogeneity.

To further investigate how RORγt^S182^ contributes to a balanced colonic Treg-Th17 program, we performed differential gene expression analysis on the Th17 subsets and LT-like Treg cells. To our surprise, the majority of RORγt^S182^-dependent genes in the two Th17 subsets were upregulated in RORγt^S182A^ cells (Figure 2A). The significant increase of *Il17a* and *Il17f* encoding the Th17 signature cytokines IL-17A and IL-17F in cluster 3 Th17 cells from the RORγt^S182A^ colon were particularly intriguing (Figure 2B-C), as RORγt is reported to function as a transcription activator for *Il17a* and *Il17f*^6^, and these results would suggest that serine 182 restricts RORγt transcription activities for a unique set of Th17 genes. In contrast to Th17 cells, RORγt^S182A^ mutant LT-like Tregs harbored a similar ratio of up and down-regulated transcripts (Figure 2A). Transcripts that were downregulated in RORγt^S182A^ cells were subset specific. For transcripts upregulated in RORγt^S182A^ cells, 9 were commonly shared among Th17 and LT-like Treg cells (Figure 2B), including the ones encoding heat shock family of proteins involved in apoptosis (Figure 2D-E). Together, these scRNA-seq results revealed that serine 182 on RORγt as the “off switch” for limiting RORγt transcription activities on select targets in colonic Th17 and LT-like Treg cells.

**Figure 2.**
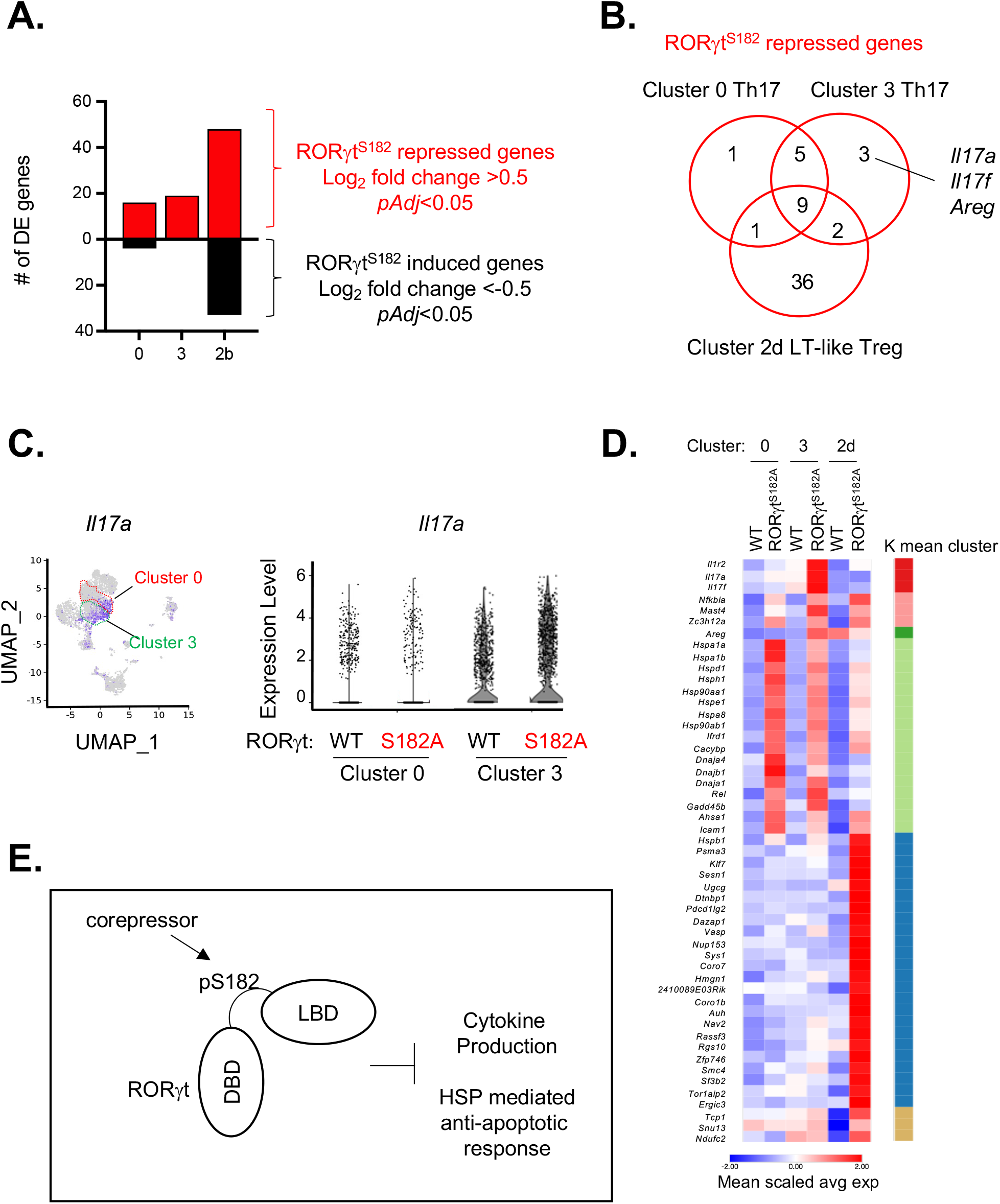
Common and distinct RORγt^S182^-dependent programs in colonic Th17 and Treg subsets. A. Number of RORγt^S182^-dependent genes in individual scRNA-seq clusters from steady state WT and RORγt^S182A^ mice (n=2). DE genes are defined as those with log_2_ fold change between RORγt^S182A^ and WT cells greater than 0.5 or less than −0.5, adjusted *p*<0.05. B. Venn diagram showing overlap and distinct RORγt^S182^-repressed genes in Th17 and Treg subsets from A. C. UMAP expression plots (left) and violin plots (right) showing expression of *Il17a* in colonic Th17 cell subsets (cluster 0 and 3) in steady state WT and RORγt^S182A^ mice. D. Heatmap of mean scaled average expression of RORγt^S182^-repressed genes from B in steady colonic Th17 (cluster 0 and 3) and LT-like Treg (cluster 2d) cells of WT and RORγt^S182A^. E. Model depicting S182 on RORγt as a negative regulator of cytokine production and heat shock response programs (HSP) in colonic T cells.

### RORγt^S182A^ augmented IL-17A production in Th17 cells in response to IL-1β stimulation

Highly homogenous Th17 cells can be generated *in vitro* from naive splenic T cells activated with anti-CD3/CD28 in the presence of IL-6 ^7^. We hypothesized that if S182 on RORγt contributed to cluster 3 Th17 cells cytokine production in a cell-intrinsic manner, we should also observe augmented IL-17 productions in Th17 cells generated from RORγt^S182A^ naïve CD4^+^ T cells in this culture system (diagramed in Figure 3A). Differential gene expression analysis revealed that cluster 3 Th17 cells expressed higher levels of *Il1r1* and *Il23r*, receptors for IL-1β and IL-23 respectively, compared to cluster 0 Th17 cells (Supplementary Figure 2C). Therefore, we assessed the IL-17 production potential in Th17 cells generated *in vitro* from WT and RORγt^S182A^ naïve CD4^+^ T cells cultured in the presence of recombinant IL-6 together with TGFβ, IL-1β, or IL-23. Interestingly, RORγt^S182A^ Th17 cells generated in the presence of IL-1β had significantly greater IL-17A and IL-17F production potential (Figure 3B and Supplementary Figure 3A). This difference is not due to altered RORγt protein abundance in the cultured WT and RORγt^S182A^ Th17 cells (Figure 3C). Western analysis confirmed that RORγt proteins were phosphorylated at S182 in cultured Th17 WT cells (Figure 3D).

**Figure 3.**
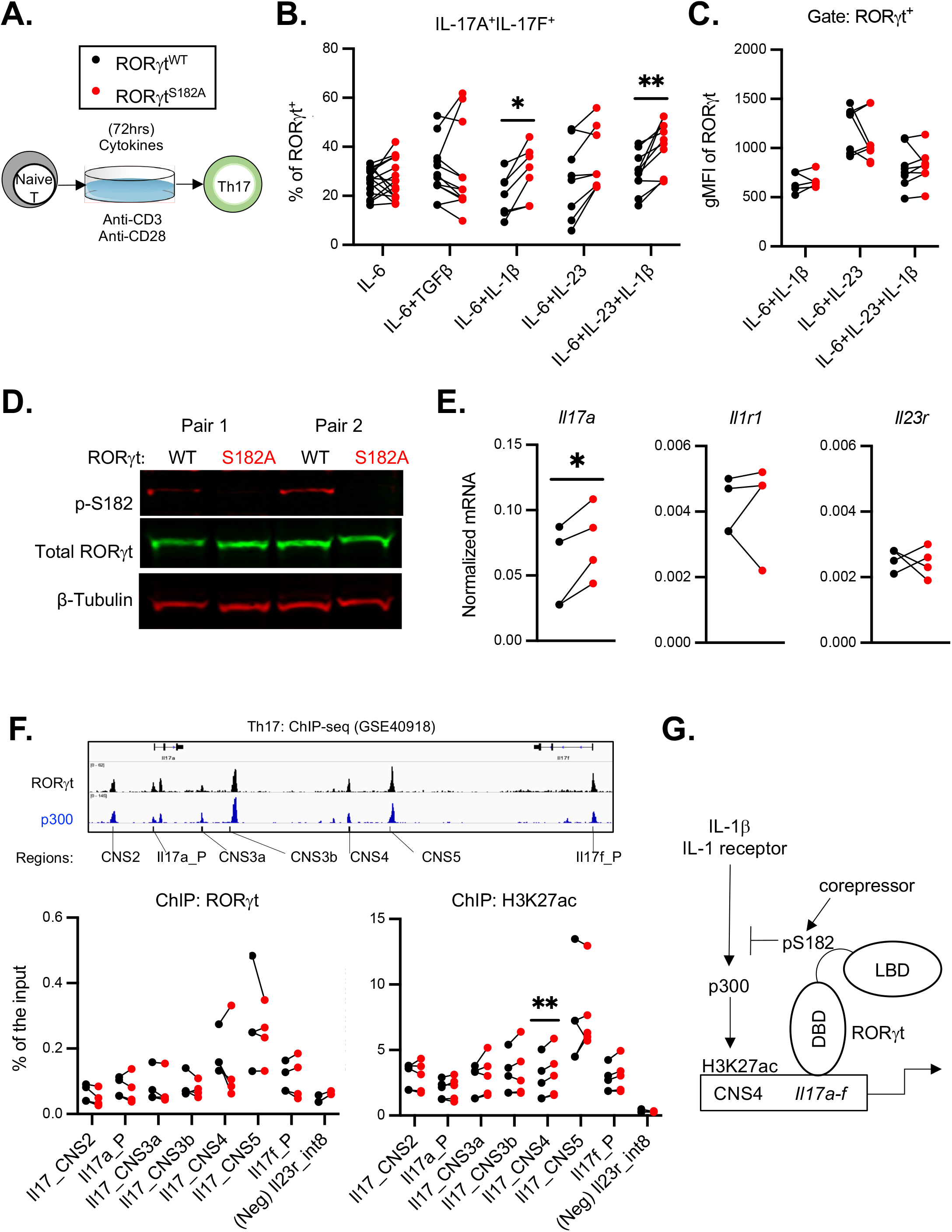
Augmented cytokine production in RORγt^S182A^ Th17 cells in response to IL-1β signaling. A. Th17 culture workflow *in vitro*. B. Proportion of IL-17A^+^IL-17F^+^ producers among Th17 cells cultured for 3 days under conditions as indicated. Each dot represents result from one mouse. * p-value<0.05, ** p-value<0.01 (paired t-test). C. Geometric mean fluorescence (gMFI) of RORγt in WT and RORγt^S182A^ Th17 cells cultured under different conditions. Each dot represents result from one mouse. D. Western blot analysis of total RORγt and S182-phosphorylated RORγt in cultured Th17 cells from two pairs of WT and RORγt^S182A^ littermates. E. Normalized *Il17a, Il1r1, and Il23r* mRNA expression detected by qRT-PCR in WT and RORγt^S182A^ Th17 cells cultured in IL-6, IL-1β, and IL-23A for 3 days. Each dot represents result from one mouse. * p-value<0.05 (paired t-test). F. Top: IGV browser showing RORγt and p300 occupancy at *Il17a-Il17f* locus as reported previously ^7^ along with the location of ChIP-qPCR primers designed to capture each of the conserved non-coding regulatory elements (CNS). Bottom: Enrichment of RORγt and H3K27ac at the *Il17a-Il17f* locus in WT and RORγt^S182A^ Th17 cells as determined by ChIP-qPCR. n=4, ** p<0.01 (t-test). G. Working model: IL-1β signaling promotes dysregulated effector cytokines production in RORγt^S182A^ Th17 cells.

RT-qPCR confirmed that an increase of *Il17a* at the RNA level in RORγt^S182A^ Th17 cells cultured in the presence of IL-6, IL-23, and IL-1β (Figure 3E). No change of *Il1r1* and *Il23r* expression were observed. Chromatin immunoprecipitation experiments indicated that RORγt binding on the *Il17a-Il17f* locus in WT and RORγt^S182A^ Th17 cells were comparable (Figure 3F). However, significantly higher H3K27 acetylation (H3K27ac) was detected at the CNS4, a putative regulatory element of the *Il17a-Il17f* locus, in RORγt^S182A^ Th17 cells (Figure 3F). Together, these results demonstrate serine 182 on RORγt negatively regulates Th17 effector cytokine production in response to IL-1β by limiting H3K27ac deposition on the *Il17a-Il17f* locus (modeled in Figure 3G).

### RORγt^S182A^ T cells drove exacerbated diseases in two models of colitis

Previous reports suggest that Th17 cells generated in the presence of IL-6, IL-23, and IL-1β contribute to autoimmune pathologies in the intestine ^30, 31^. Therefore, we hypothesized that RORγt^S182A^ Th17 cells with hyperresponsiveness to IL-1β stimulation would have the potential to exacerbate tissue inflammation. To test this possibility in a model of T cell transfer colitis, WT or RORγt^S182A^ CD4 naive T cells were introduced into RAG1^-^ recipients lacking endogenous T cells. WT and RORγt^S182A^ T cell recipients all experienced similar weight loss between day 2-16. However, the weights of WT recipients stabilized between day 19-31, but the weights of those with RORγt^S182A^ cells were further reduced (Figure 4A). RT-qPCR confirmed a significant increase of *Il17a* mRNA were found in the colonic lamina propria cells from the RORγt^S182A^ recipients (Figure 4B). These results indicate that S182 on RORγt and/or its phosphorylation are negative regulators of T cell mediated colonic inflammation.

**Figure 4.**
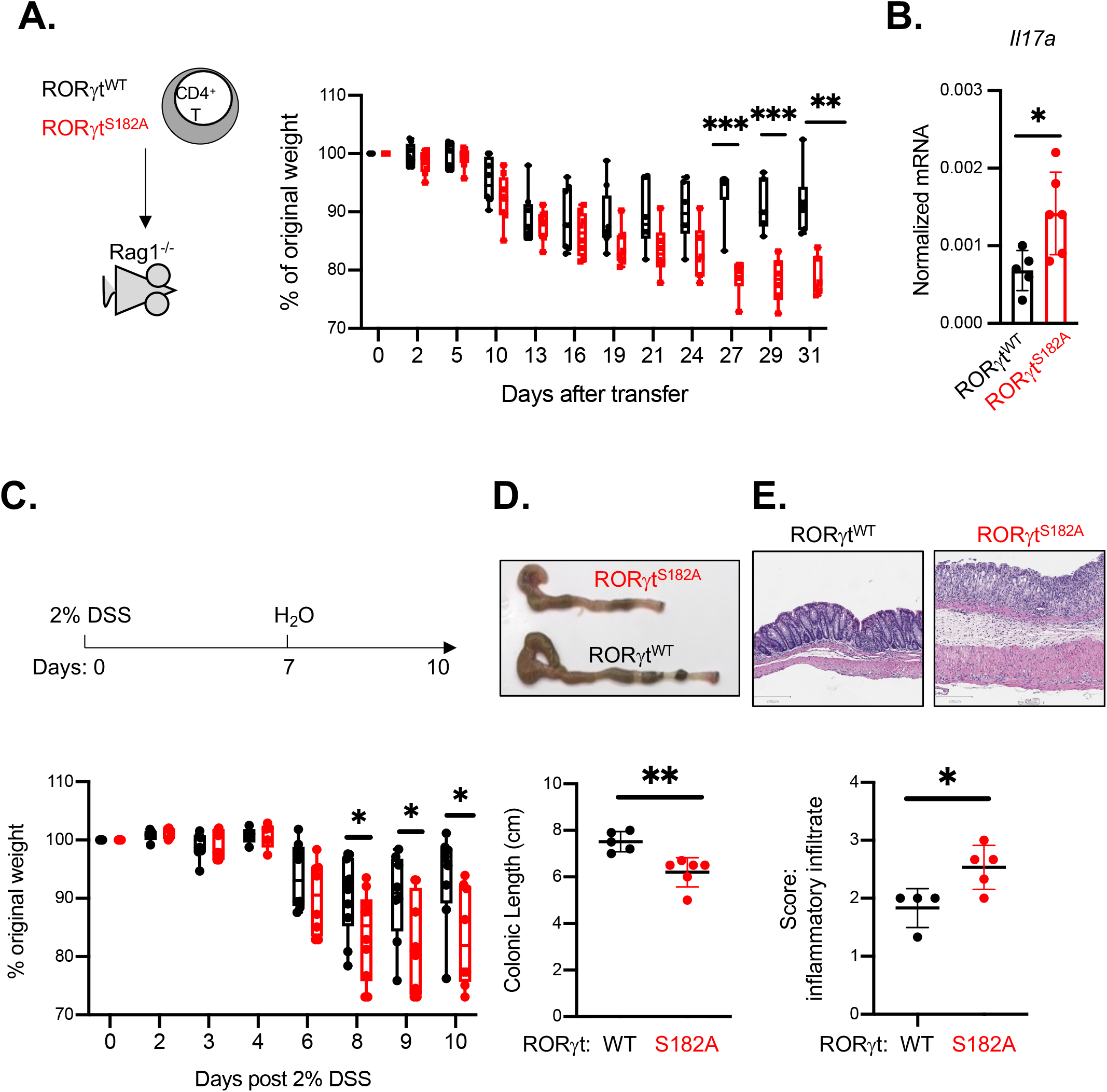
RORγt^S182A^ T cells drove exacerbated diseases in two models of colitis. A. Weight changes of Rag1^-/-^ mice receiving WT or RORγt^S182A^ naive CD4^+^ T cells. Each dot represents result from one mouse. ** p-value<0.01 and *** p-value<0.001 (multiple t-test). B. Expression level of *Il17a* in colonic lamina propria of Rag1^-/-^ mice receiving WT or RORγt naive CD4^+^ T cells. Each dot represents result from one mouse. * p-value<0.05 (t-test). C. Weight changes of WT and RORγt^S182A^ mice challenged with 2% DSS in drinking water for 7 days and monitored for another 3 days. Each dot represents result from one mouse. * p-value<0.05 (multiple t-test). D. Top: representative bright field images of colons from C. Bottom: summarized colonic lengths of DSS treated WT and RORγt^S182A^ mice from C harvested on day 10. Each dot represents result from one mouse. ** p-value<0.01 (t-test). E. Top: representative colonic sections from D. Bottom: summarized score of colonic inflammatory infiltrates (bottom) in DSS treated WT and RORγt^S182A^ mice. Each dot represents result from one mouse. * p-value<0.05 (t-test).

In addition, we also assessed the contribution of RORγt^S182^ in DSS induced acute intestine epithelial injury model to yield disease pathologies similar to those observed in human ulcerative colitis patients ^32^. Similar to our findings in the T cell transfer colitis model, DSS challenged RORγt^S182A^ mice had a significant delay in weight recovery on day 8-10 as compared to their WT littermates (Figure 4C). Colons harvested from DSS-challenged RORγt^S182A^ mice were much shorter than those from DSS-challenged WT mice (Figure 4D). Histological analysis revealed significantly more infiltrating immune cells present in the colons from RORγt^S182A^ mice day 10 post DSS challenge (Figure 4E). These *in vivo* results suggest that S182 on RORγt and/or its phosphorylation in CD4^+^ T cells contribute to inflammation resolution in colitis settings.

### scRNA-seq revealed RORγt^S182^-dependent colonic Th17 and Treg programs during DSS challenge

To characterize the contribution of RORγt^S182A^ in T cell programs during the resolution phase of colonic inflammation, we harvested colonic lamina propria cells from two pairs of WT and RORγt^S182A^ mice on day 10 post DSS challenge for scRNA-seq. In WT mice, DSS challenge increased the proportion of colonic memory (cluster 1) and Treg (cluster 2) cells, as well as, altered cluster 0 and 3 Th17 subset ratio to 2:1 (Supplementary Figure 4A-B). DSS-responsive genes in cluster 0 and 3 Th17 cells (Supplementary Figure 4C) were enriched with tumor necrosis factor-alpha (TNF) and oxidative phosphorylation signatures (Supplementary Figure 4D), which are pivotal pathways previously implicated in colitis ^33, 34^ Compared to WT tissues, DSS-challenged RORγt^S182A^ colons harbored fewer memory T cells (cluster 1) (Supplementary Figure 5A).

Differential gene expression analysis revealed greater number of genes upregulated in the DSS-challenge RORγt^S182A^ colonic Th17 and Treg subsets (Supplementary Figure 5B). This confirmed the repressive role of S182 on RORγt in regulating colonic T cell programs in inflamed settings, similar to what we observed under steady state (Figure 2A). Interestingly, while *Il17a* expression in cluster 3 Th17 cells from the DSS treated mice was no longer subject to regulation by RORγt^S182^, *Il17a* expression in cluster 0 Th17 cells became sensitive to RORγt^S182^ regulation. Cluster 0 Th17 cells from DSS treated RORγt^S182A^ mice had higher *Il17a* expression (Figure 5A-B). Flow cytometry analysis confirmed that the RORγt^+^Foxp3^-^ Th17 cells had more IL-17A production potential in the DSS treated RORγt^S182A^ colon (Figure 5C).

**Figure 5.**
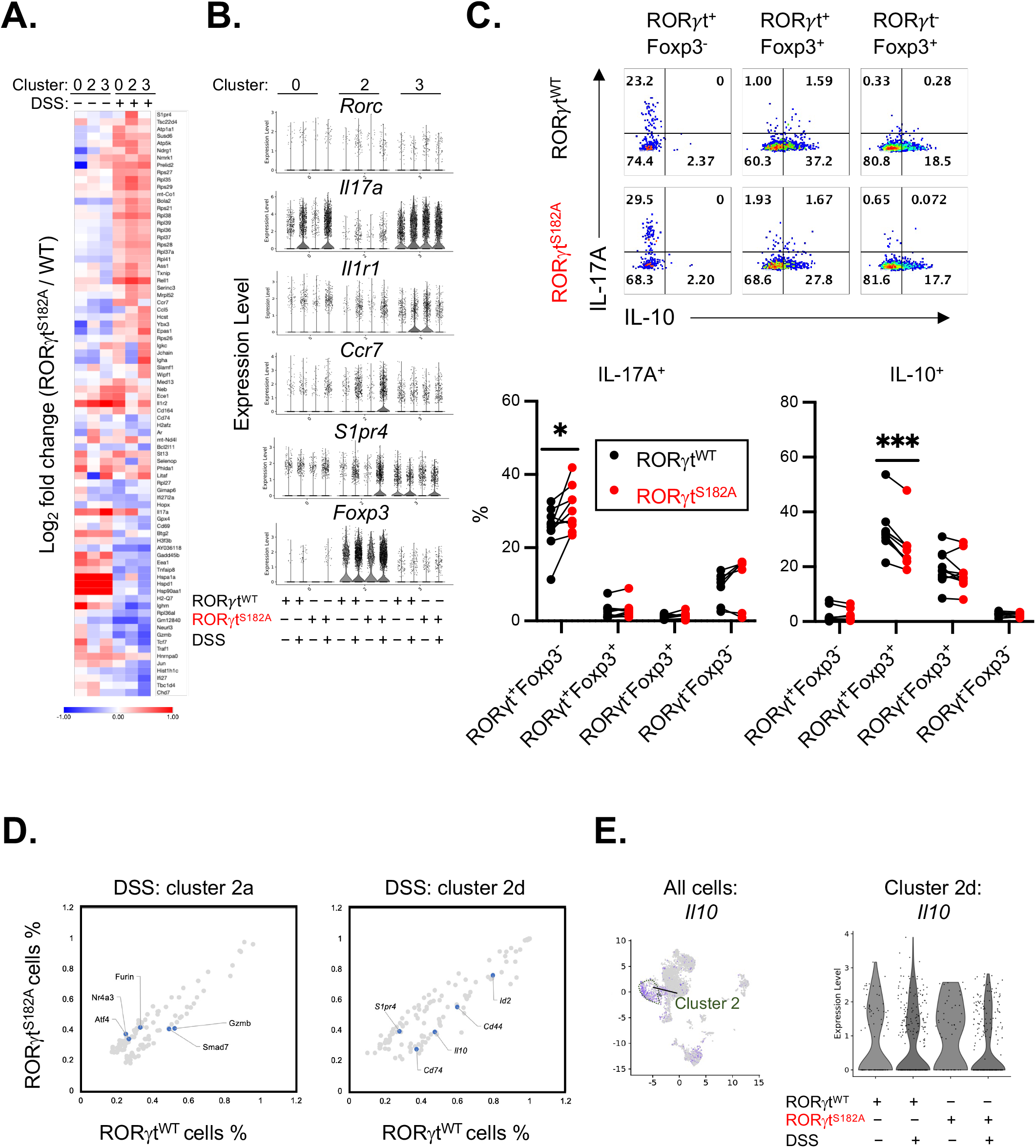
RORγt^S182A^-dependent colonic Th17 and Treg programs during DSS challenge. A. Heatmap of Log_2_ fold change of select RORγt^S182^-dependent genes expression in Th17 (cluster 0 and 3) and Treg (cluster 2) from steady state and DSS challenged colons. B. Violin plots of the selected gene expressions in Th17 (cluster 0 and 3) and Treg (cluster 2) cells. C. Top: representative flow cytometry analysis. Bottom: proportion of IL-10 and IL-17A production potential in colonic RORγt^+^Foxp3^-/-^ (Th17), RORγt^+^Foxp3^+^ (RORγt^+^ Treg), RORγt^-^Foxp3^+^ (Treg), and RORγt^-^Foxp3^-^ cells from DSS treated WT and RORγt^S182A^ mice and harvested on day 10. Each dot represents result from one mouse. * p-value<0.05, *** p-value<0.001 (t-test). D. Percentage of cells expressing RORγt^S182^-dependent genes (grey, adjusted *p*<0.05) in colonic Treg subsets 2a and 2d from DSS challenged WT and RORγt^S182A^ mice. Select genes were labeled and highlighted in blue. E. Left: UMAP displaying *Il10* expression in scRNA-seq dataset. Right: violin plots of the *Il10* expression in LT-like Treg (subset 2d) cells from control or DSS challenged WT and RORγt^S182A^ mice.

In addition to the augmented IL-17A production potential in Th17 cells, DSS-challenged RORγt mice also had significantly fewer IL-10 producers among the RORγt Treg cells (Figure 5C), which was reported to be key negative regulators of colonic inflammation ^35–38^. scRNA-seq analysis revealed that RORγt^S182^ regulated *Il10* expression in LT-like (subset 2d) Tregs in the DSS challenged colon (Figure 5D-E). Altogether, these data indicate that RORγt^S182^ negatively regulates effector cytokine production in Th17 cells and promotes expression of anti-inflammation cytokine IL-10 in RORγt^+^ Treg cells to facilitate inflammation resolution during colitis (modeled in Supplementary Figure 6A).

### RORγt^S182A^ mice experienced the more severe disease in EAE

Finally, we assessed whether S182 of RORγt also negatively regulates inflammation outside of the intestine. Previous reports suggest that pathogenic Th17 cells promote disease in the EAE mouse model ^39^ by secreting IL-17A to recruit CD11c^+^ dendritic cells and driving additional IL-17A production from bystander TCRγδ^+^ type 17 (Tγδ17) cells to fuel local inflammation ^6, 40–42^ as modeled in Figure 6A. When challenged in this model, RORγt^S182A^ mice had higher disease scores and experienced greater weight loss (Figure 6B-C). Total spinal cord immune infiltrates from MOG immunized RORγt^S182A^ mice had significantly higher levels of *Cd4, Cd11c*, and *Il17a* mRNAs (Figure 6D-E). Flow cytometry results also confirmed a significant increase in the number of IL-17A producing Th17 (CD4^+^TCRγδ^-^) and Tγδ17 (CD4^-^TCRγδ^+^) cells (Figure 6F). While greater number of infiltrated DC (CD11c^+^) were found in the spinal cord of the MOG immunized RORγt^S182A^ mice, no change in macrophages (CD11b^+^F4/80^+^) were observed (Figure 6G). Overall, these results demonstrate the critical role of S182 on RORγt in modulating T cell functions to protect against colonic mucosal inflammation and central nervous system autoimmune pathologies.

**Figure 6.**
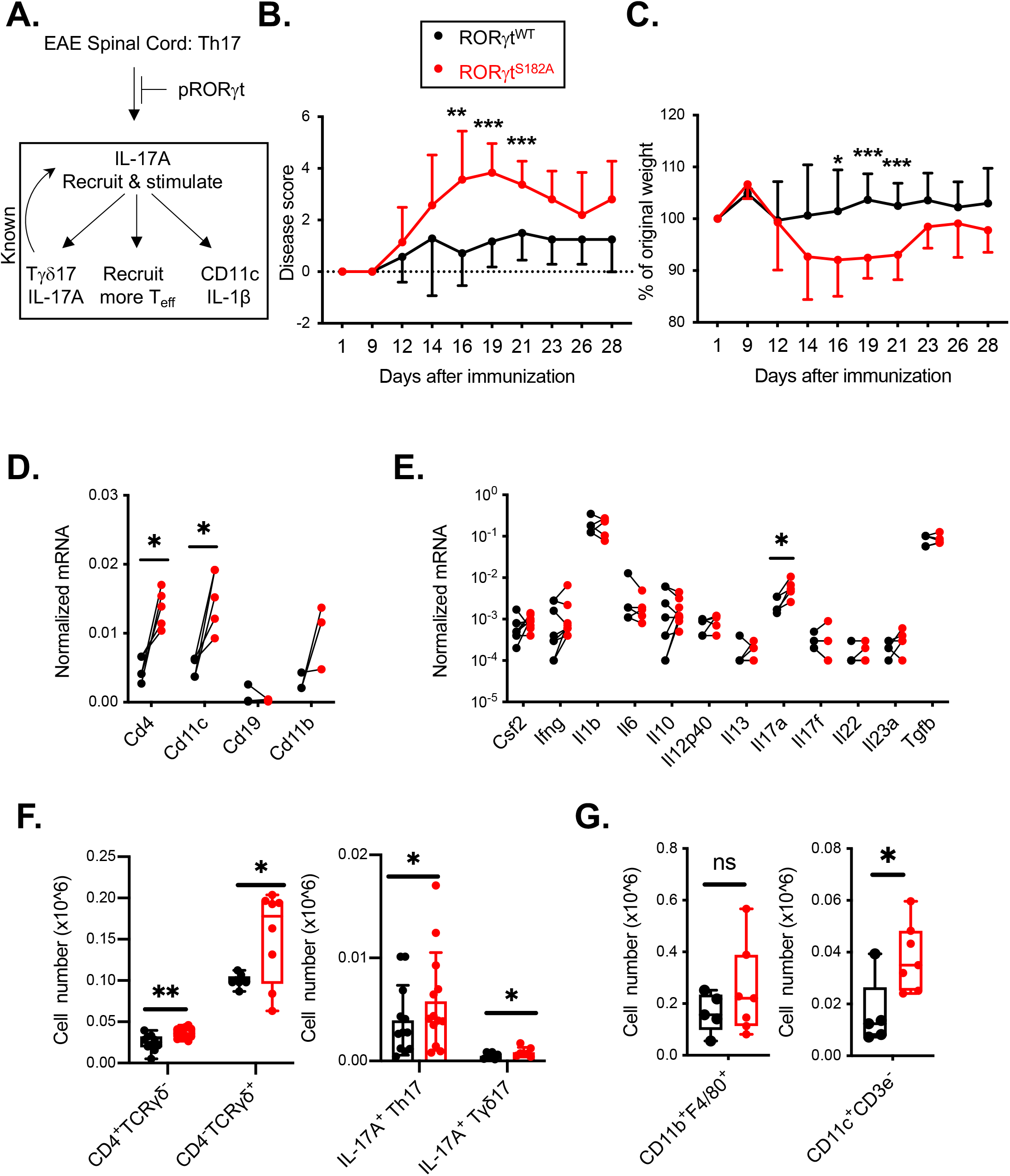
RORγt^S182A^ mice challenged in the EAE model experienced more severe disease. A. Working hypothesis: RORγt^S182^ acts as the upstream negative regulator of pathogenic Th17 cell activities in EAE. B. Disease score of MOG-immunized WT (n=7) and RORγt^S182A^ (n=14) mice. Results were combined of three independent experiments. ** p-value<0.01, *** p-value<0.001 (multiple t-test). C. Weight change of MOG-immunized WT and RORγt^S182A^ mice. Results were combined of three independent experiments. * p-value<0.05, *** p-value<0.001 (multiple t-test). D. Normalized mRNA expression of *Cd4, Cd11c, Cd19* and *Cd11b* in spinal cords infiltrated at the peak of EAE disease (day 16-19) from MOG-immunized WT and RORγt^S182A^ mice as detected by qRT-PCR. Each dot represents result from one mouse. * p-value<0.05 (paired t-test). E. Normalized mRNA expression of the indicated cytokine genes in spinal cords infiltrated at the peak of EAE disease (day16-19) from MOG-immunized WT and RORγt^S182A^ EAE mice as detected by qRT-PCR. Each dot represents result from one mouse. * p-value<0.05 (paired t-test). F. Number of total T helper (CD4^+^) and TCRγδ^+^ (left) and IL-17A^+^ expressing Th17 and Tγδ17 cells (right) in EAE mice harvested at the peak of disease (day16-19). Each dot represents result from one mouse. * p-value<0.05 (t-test). G. CD11b^+^F4/80^+^ (macrophage, left) and CD3ε^-^CD11c^+^ (DC, right) cells numbers in the spinal cords of EAE mice harvested at the peak of disease (day16-19). Each dot represents result from one mouse. * p-value<0.05 (t-test).

## DISCUSSION

Our single-cell transcriptomic studies in lamina propria CD4^+^ T cells identified two subsets of Th17 with distinct cell surface receptors and effector molecule expressions in the steady state colon. Colonic cluster 0 Th17 cells express abundant transcripts encoding various chemokines, including CCL2, CCL5, CCL7, CCL8, CXCL11, and CXCL13, suggesting that they may be involved in orchestrating chemotactic gradients that regulate local immune cells trafficking. Colonic cluster 3 Th17 cells express higher levels of transcripts encoding receptors for local inflammation signals, including IL-1 and IL-23 receptors. The generation of Th17 cells and their effector functions depend on the transcription activities of RORγt ^5–7^. However, it is unclear how RORγt contributes to distinct functions of different Th17 subsets. In this report, we showed that RORγt is phosphorylated at S182 in Th17 cells. Although this residue is dispensable for Th17 differentiation, it is essential for maintaining the 1:1 balance ratio of cluster 0 and cluster 3 Th17 cells in the steady state colon.

Single-cell transcriptomic studies revealed that serine 182 negatively regulates RORγt-dependent Th17 gene programs in a context specific manner. Under steady state, S182 on RORγt restricts IL-17 production in cluster 3 Th17 cells. During DSS-induced colitis, S182 on RORγt restricts IL-17 production in cluster 0 Th17 cells instead. The molecular mechanism underlying this context-dependent regulation remains to be elucidated. Results from *in vitro* Th17 culture studies suggest that S182 on RORγt restricts H3K27 acetylation at the *Il17a-Il17f* locus to prevent exacerbated Th17 cytokine production when cells were stimulated by the inflammatory cytokine IL-1β. Other signaling pathways downstream of IL-6, TGFβ, and/or IL-23 failed to engage the S182 regulatory node. These results suggest that alterations of stimulatory cytokines in the microenvironment may underly the dynamic switch of cluster 0 and 3 Th17 cells’ sensitivity toward S182 regulation during DSS-induced colitis.

RORγt is also expressed in subsets of Tregs abundantly found in the intestinal mucosa and peripheral lymphoid organs ^35, 36^. Secretion of IL-10 by these cells helps to protect against autoimmune diseases ^16, 35, 36, 38, 43^ by dampening uncontrolled production of inflammatory cytokines, including IL-17A, in mouse models ^44^. Reduced IL-10 level is associated with severe cases of inflammatory bowel diseases in human patients ^45, 46^. Administration of IL-10 has been proposed as a potential therapy for these patients ^45, 46^. In this study, we revealed that S182 on RORγt is required for maintaining LT-like (cluster 2d) Treg population in the steady state colon and their IL-10 production potential during DSS-induced colitis. In addition to a loss of IL-10 expression, RORγt^S182A^ mutant LT-like Tregs had decreased levels of *Cd44* and *Id2*, encoding molecules implicated in Treg activation and IL-10 production ^47, 48^, and increased expression of *S1pr4*, which is involved in cell migration ^*49*^.

The imbalance of Th17-Treg populations and functions in RORγt^S182A^ mutant mice resulted in delayed recovery and more pathology when challenged in the colitis and EAE models. These findings highlight the previously unappreciated role of the RORγt hinge region harboring S182 for keeping the Th17 programs in check. Future studies will be needed to investigate whether RORγt^S182^-dependent LT-like Tregs and Th17 subsets exert paracrine effects on other local immune cell programs to drive the delayed DSS and EAE disease recovery phenotypes observed in RORγt^S182A^ mice. Furthermore, it remained to be explored whether the same or distinct kinase(s) are involved in phosphorylating S182 on RORγt in different T cell subsets. Future proteomics studies will be needed to uncover the corepressor and/or coactivator complexes interacting with the hinge region of RORγt in a S182 dependent manner.

Given the essential role of RORγt in the differentiation and functions of diverse T cell subsets implicated in autoimmunity, many RORγt inhibitors have been developed for therapeutic purposes. Most current inhibitors are designed to disrupt RORγt-dependent transcription by preventing its interaction with steroid receptor coactivators. Unfortunately, as this is a common mechanism RORγt employs across the different cell and tissue types, this approach will likely exert undesired side-effects on thymic development and other innate immune cells expressing RORγt. Our discovery of the involvement of S182 on RORγt in regulating Th17 and Treg functions, but the dispensable role in T cell development, provides a unique venue for designing better therapies to combat T cell-mediated inflammatory diseases.

## METHODS

### Mice

RORγt^S182A^ were generated by CRISPR-Cas9 technology in the C57BL/6 (Jackson Laboratories) background mice and confirmed by sanger sequencing of the *Rorc* locus. Heterozygous mice were bred to yield 8-12 WT and homogenous knock-in (RORγt^S182A^) cohoused littermates for paired experiments. Rag1^-/-^’ (Jackson Laboratory stock No: 002216) were obtained from Dr. John Chang’s Laboratory. Adult mice at least eight weeks old were used. All animal studies were approved and followed the Institutional Animal Care and Use Guidelines of the University of California San Diego.

### Th17 cell culture

Mouse naive T cells were purified from spleens and lymph nodes of 8-12 weeks old mice using the Naive CD4^+^ T Cell Isolation Kit according to the manufacturer’s instructions (Miltenyi Biotec). Cells were cultured in Iscove’s Modified Dulbecco’s Medium (IMDM, Sigma Aldrich) supplemented with 10% heat-inactivated FBS (Peak Serum), 50U/50 ug penicillin-streptomycin (Life Technologies), 2 mM glutamine (Life Technologies), and 50 μM β-mercaptoethanol (Sigma Aldrich). For polarized Th17 cell polarization, naive cells were seeded in 24-well or 96-well plates, pre-coated with rabbit anti-hamster IgG, and cultured in the presence of 0.25 μg/mL anti-CD3ε (eBioscience), 1 μg/mL anti-CD28 (eBioscience), 20 ng/mL IL-6 (R&D Systems), and/or 0.1 ng/mL TGF-β (R&D Systems), 20 ng/mL IL-1β (R&D Systems), 25 ng/mL IL-23 (R&D Systems) for 72 hours.

### DSS induced and T cell transfer colitis

Dextran Sulfate Sodium Salt (DSS) Colitis Grade 36,000-59,000MW (MP Biomedicals) was added to the drinking water at a final concentration of 2% (wt/vol) and administered for 7 days. Mice were weighed every other day. On day 10, colons were collected for H&E staining and lamina propria cells were isolation as described ^20^. Cells were kept for RNA isolation or flow cytometry. The H&E slides from each sample were scored in a double-blind fashion as described previously ^21^. For T cell transfer model of colitis, 0.5 million naive CD4^+^ T cells isolated from mouse splenocytes using the Naive CD4^+^ T Cell Isolation Kit (Miltenyi), as described above, were injected intraperitoneally into RAG1^-/-^ recipients. Mice weights were measured twice a week. Pathology scoring of distal colons from DSS-challenged mice were performed blind following previously published guidelines ^22^.

### scRNA-seq and analysis

Colonic lamina propria cells from DSS or non-DSS treated mice were collected and enriched for CD4^+^ T cells using the mouse CD4^+^ T cell Isolation Kit (Miltenyi). Enriched CD4^+^ cells (~10,000 per mouse) were prepared for single cell libraries using the Chromium Single Cell 3’ Reagent Kit (10xGenomics). The pooled libraries of each sample (20,000 reads/cell) were sequenced on one lane of NovaSeq S4 following manufacturer’s recommendations.

Cellranger v3.1.0 was used to filter, align, and count reads mapped to the mouse reference genome (mm10-3.0.0). The Unique Molecular Identifiers (UMI) count matrix obtained was used for downstream analysis using Seurat (v4.0.1). The cells with mitochondrial counts >5%, as well as outlier cells in the top and bottom 0.2% of the total gene number detected index were excluded. After filtering, randomly selected 10,000 cells per sample were chosen for downstream analysis. Cells with *Cd4* expression lower than 0.4 were removed, resulting in 27,420 total cells from eight samples. These cells were scaled and normalized using log-transformation, and the top 3,000 genes were selected for principal component analysis. The dimensions determined from PCA were used for clustering and non-linear dimensional reduction visualizations (UMAP). Differentially expressed genes identified by FindMarkers were used to characterize each cell cluster. Other visualization methods from Seurat such as VlnPlot, FeaturePlot, and DimPlot were also used.

### EAE model

EAE was induced in 8-week-old mice by subcutaneous immunization with 100 μg myelin oligodendrocyte glycoprotein (MOG35-55) peptide (GenScript Biotech) emulsified in complete Freund’s adjuvant (CFA, Sigma-Aldrich), followed by administration of 400 ng pertussis toxin (PTX, Sigma-Aldrich) on days 0 and 2 as described ^23^. Clinical signs of EAE were assessed as follows: 0, no clinical signs; 1, partially limp tail; 2, paralyzed tail; 3, hind limb paresis; 4, one hind limb paralyzed; 5, both hind limbs paralyzed; 6, hind limbs paralyzed, weakness in forelimbs; 7, hind limbs paralyzed, one forelimb paralyzed; 8, hind limb paralyzed, both forelimbs paralyzed; 9, moribund; 10, death.

### Flow cytometry

Cells were stimulated with 5 ng/mL Phorbol 12-myristate 13-acetate (PMA, Millipore Sigma) and 500ng/mL ionomycin (Millipore Sigma) in the presence of GoligiStop (BD Bioscience) for 5 hours at 37°C, followed by cell surface marker staining. Fixation/Permeabilization buffers (eBioscience) were used per manufacturer instructions to assess intracellular transcription factor and cytokine expression. Antibodies are listed in Supplementary Table 1.

### cDNA synthesis, qRT-PCR, and RT-PCR

Total RNA was extracted with the RNeasy kit (QIAGEN) and reverse transcribed using iScript™ Select cDNA Synthesis Kit (Bio-Rad Laboratories). Real time RT-PCR was performed using iTaq™ Universal SYBR^®^ Green Supermix (Bio-Rad Laboratories). Expression data was normalized to *Gapdh* mRNA levels. Primer sequences are listed in Supplementary Table 2.

### Chromatin immunoprecipitation

ChIP was performed on 5-10 million Th17 cells crosslinked with 1% formaldehyde. Chromatin was sonicated and immunoprecipitated using antibodies listed in Table S1 and captured on Dynabeads (ThermoFisher Scientific). Immunoprecipitated protein-DNA complexes were reverse cross-linked and chromatin DNA purified as described ^24^. Primer sequences are listed in Supplementary Table 2.

### Statistical analysis

All values are presented as means ± SD. Significant differences were evaluated using GraphPad Prism 8 software. The student’s t-test or paired t-test were used to determine significant differences between two groups. A two-tailed p-value of <0.05 was considered statistically significant in all experiments.

## AUTHOR CONTRIBUTIONS

S.M. designed and performed *in vivo, in vitro* and scRNA-seq studies with the help of N.C. scRNA-seq study was analyzed by S.P. with input from J.T.C and W.J.M.H. S.M. and B.S.C. completed the DSS colitis studies. P.R.P. completed the double-blinded histology scoring of the colonic sections from DSS challenged mice. N.A. performed the western analysis. J.T.C. supervised the scRNA-seq analysis by S.P., contributed resources, and edited the manuscript. W.J.M.H. wrote the manuscript together with S.M.

## ACNKOWLEDGEMENTS

S.M., N.C., B.S.C., P.R.P, N.A. and W.J.M.H. were partially funded by the Edward Mallinckrodt, Jr. Foundation and the National Institutes of Health (NIH) (R01 GM124494 to WJM Huang). S.P. and J.T.C. were supported by National Institutes of Health grant AI123202 and AI132122 (JTC). Illumina sequencing was conducted at the IGM Genomics Center, University of California San Diego, with support from NIH (S10 OD026929). The Moores Cancer Center Histology Core conducted colonic tissue sectioning and staining with support from NIH (P30 CA23100). We thank Karen Sykes for suggestions to the manuscript.

