## Supplemental Figure S1-6 and Supplemental Table S1-2 for "RORγt serine 182 tightly regulates T cell heterogeneity to maintain mucosal homeostasis and restrict tissue inflammation"

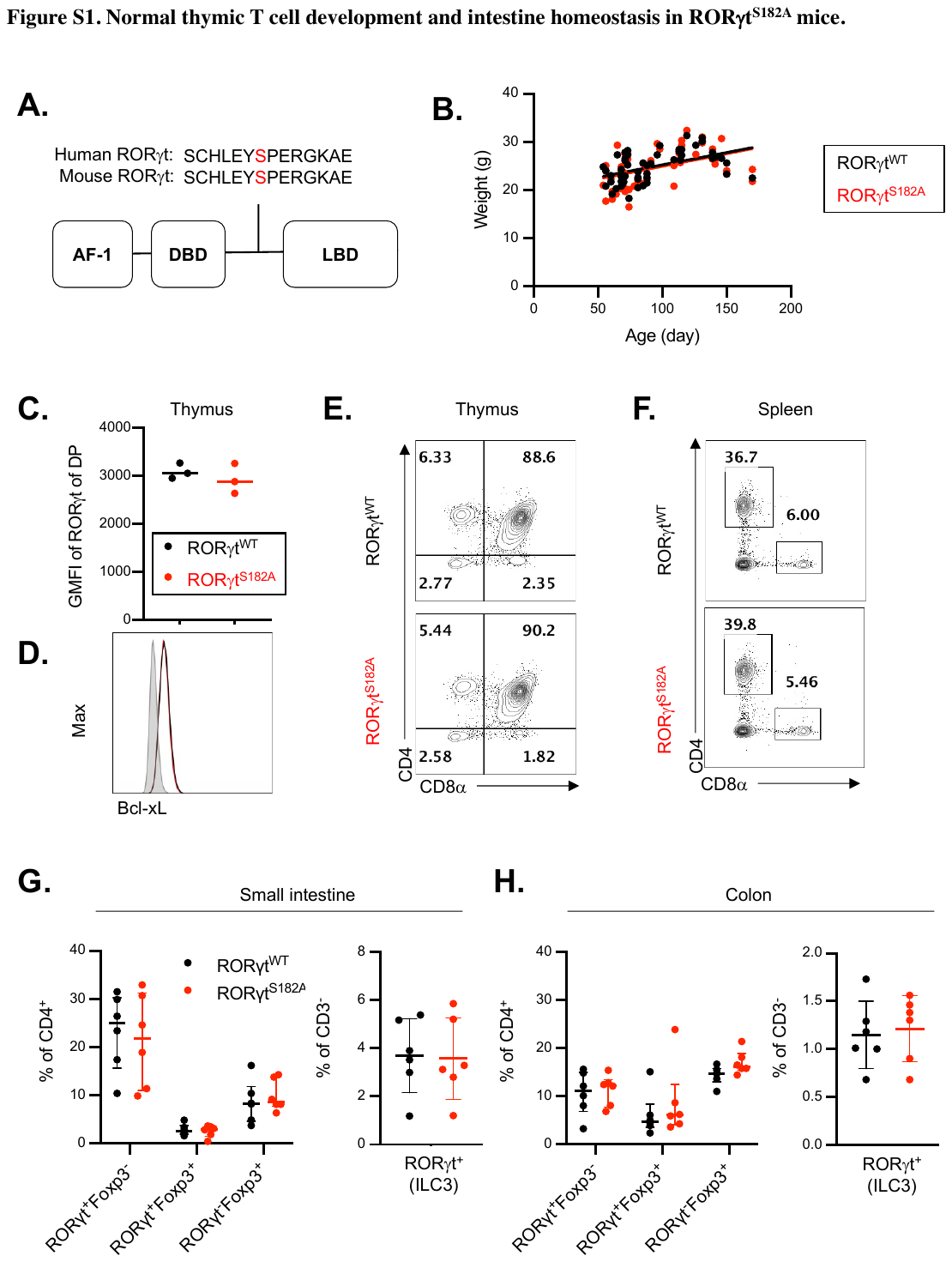

**Supplementary** **Figure 1. Normal thymic T cell development and intestine homeostasis in RORγt^S182A^ mice.**

A. S182 (black line) of RORγt is conserved between mouse and human. AF-1, Activation Function; DBD, DNA binding domain; LBD, ligand binding domain.
B. Weight of WT (n=49) and RORγt^S182A^ (n=57) mice assessed at the indicated ages. Each dot represents result from one mouse.
C. Geometric mean fluorescent intensity (GMFI) of RORγt in thymic CD4^+^CD8α^+^ double positive (DP) cells from WT and RORγt^S182A^ cohoused littermates. Each dot represents result from one mouse.
D. Representative histogram of Bcl-xL expression in thymic DP cells from WT and RORγt^S182A^ mice. This experiment was repeated three times on independent biological samples with similar results.
E. Representative flow cytometry analysis of CD4 and CD8a expression in thymic cells from WT and RORγt^S182A^ mice. This experiment was repeated three times on independent biological samples with similar results.
F. Representative flow cytometry analysis of CD4 and CD8α expression in splenocytes from WT and RORγt^S182A^ mice. This experiment was repeated three times on independent biological samples with similar results.
G. Proportion of RORγt^+^Foxp3^-^, RORγt^+^Foxp3^+^, and RORγt^-^Foxp3^+^ T cell subsets and CD3e^-^RORγt^+^Foxp3^-^ (ILC3) cells in the steady state small intestinal lamina propria of WT (n=6) and RORγt^S182A^ mice (n=6).
H. Proportion of RORγt^+^Foxp3^-^, RORγt^+^Foxp3^+^, and RORγt^-^Foxp3^+^ T cell subsets and CD3e^-^RORγt^+^Foxp3^-^ ILC3 cells in the steady state colonic lamina propria of WT (n=6) and RORγt^S182A^ mice (n=6).

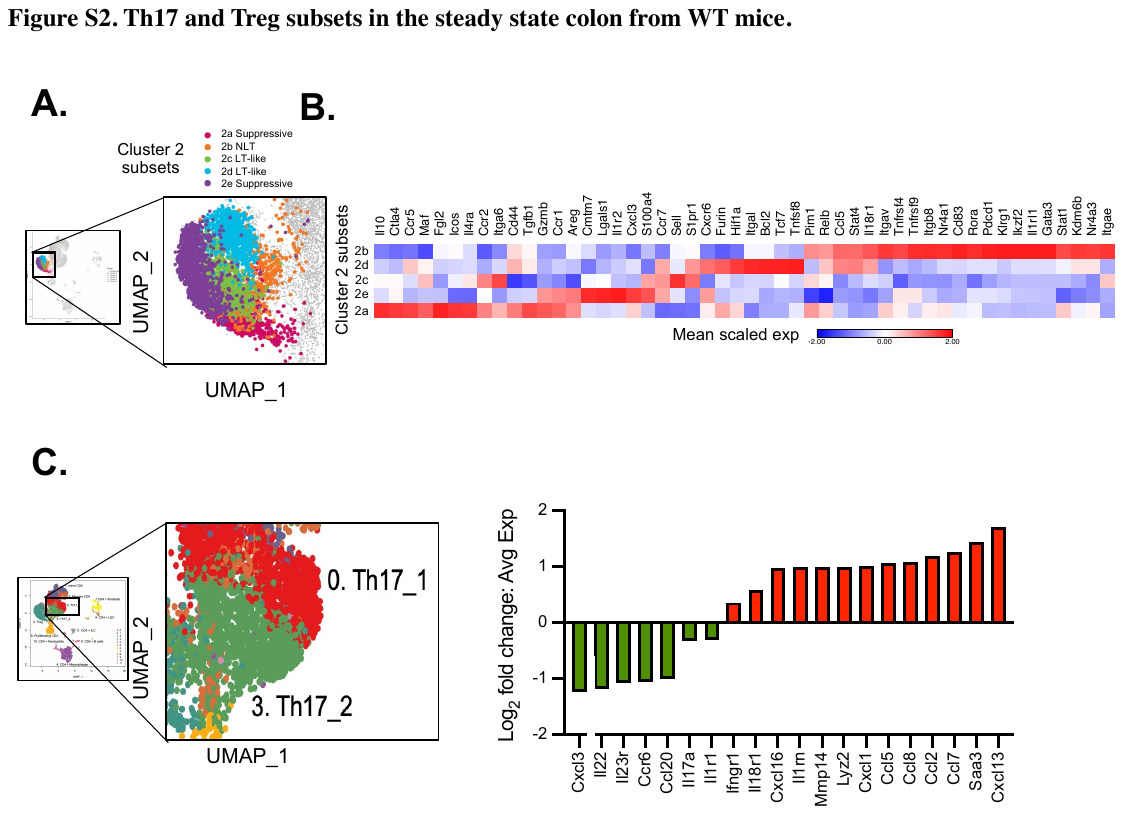

**Supplementary** **Figure 2. Th17 and Treg subsets in the steady state colon from WT mice.**

A. Closed up UMAP of the five Treg subsets in Cluster 2.
B. Heatmap of mean scaled average expression of select Treg subsets enriched genes from A.
C. Left: Closed up UMAP plot showing two neighboring populations of colonic Th17 cells in red (cluster 0) and green (cluster 3). Right: select cell surface receptors and effector molecules enriched in the two Th17 subsets from WT mice.

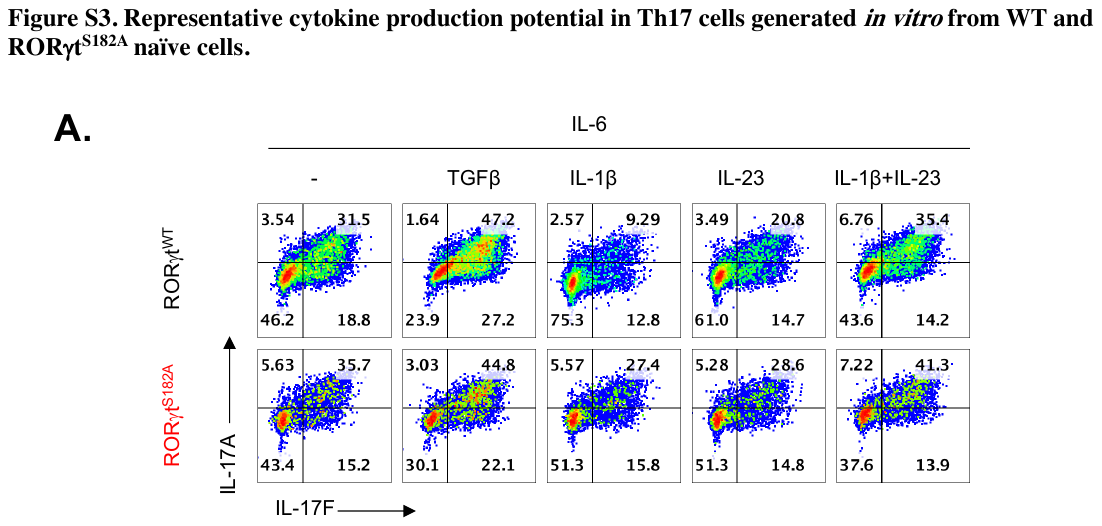

**Supplementary** **Figure 3**. **Representative cytokine production potential in Th17 cells generated *in vitro* from WT and RORγt^S182A^ naïve cells.**

A. Representative flow cytometry analysis of IL-17A and IL-17F expression in cultured Th17 cells from WT and RORγt^S182A^ mice as described in Figure 3B.

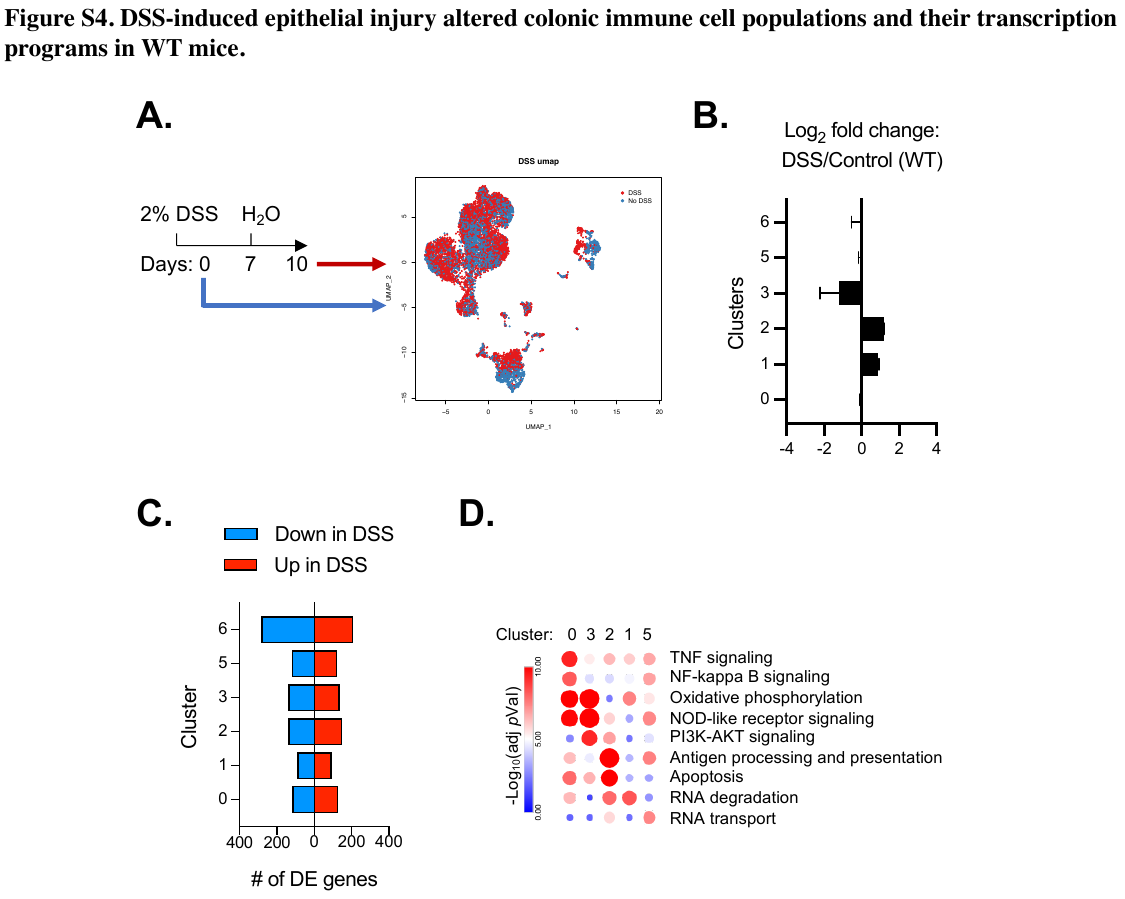

**Supplementary** **Figure 4. DSS-induced epithelial injury altered colonic immune cell populations and their transcription programs in WT mice.**

A. UMAP plot of colonic lamina propria cells obtained from steady state (blue) and day 10 post DSS challenged (red) WT mice.
B. Altered proportions of each CD4^+^ T cell cluster in colonic lamina propria cells from steady state (n=2) and day 10 post DSS treated WT mice (n=2).
C. DSS-dependent genes (*p* <0.05) identified in each cell cluster.
D. Heatmap of -Log_10_ *p*-Value of the top seven KEGG pathways from DSS-dependent genes in clusters 0,1, 2, 3, and 5.

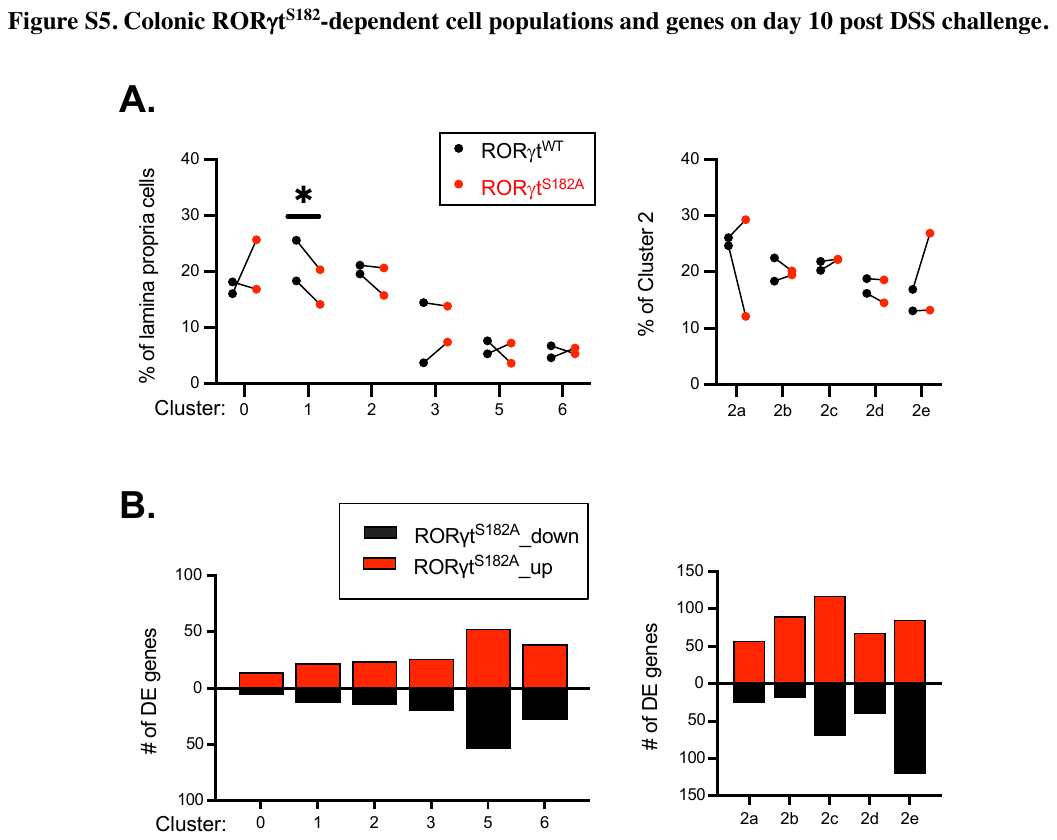

**Supplementary** **Figure 5. Colonic RORγt^S182^-dependent cell populations and genes on day 10 post DSS challenge.**

A. Proportions of individual scRNA-seq clusters among colonic lamina propria CD4^+^ T cells from DSS-challenged WT and RORγt^S182A^ mice. * p-value<0.05, n=2.
B. Number of RORγt^S182^-dependent genes in individual scRNA-seq clusters from DSS-challenged WT and RORγt^S182A^ mice (n=2). # of DE genes means number of differentially expressed genes.

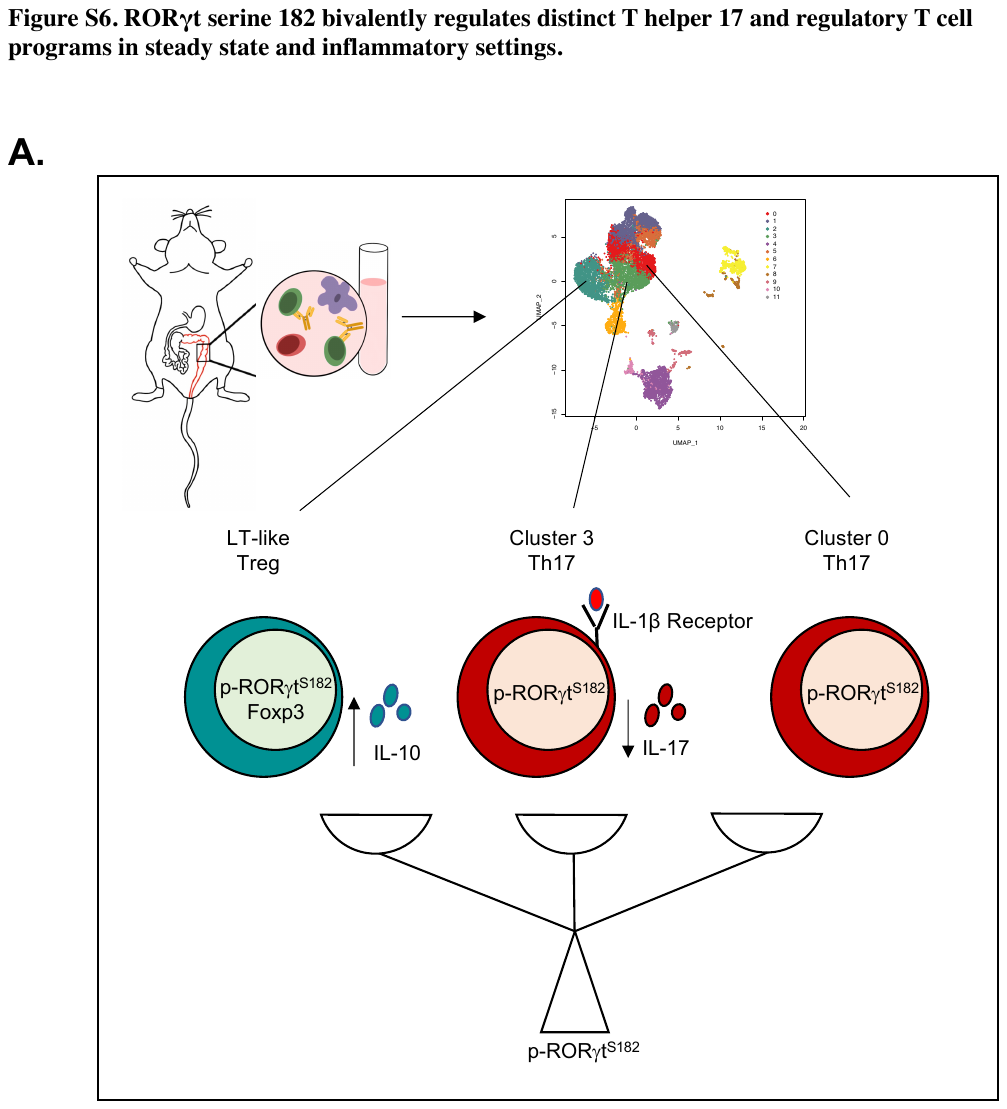

**Supplementary** **Figure 6. RORγt serine 182 bivalently regulates distinct T helper 17 and regulatory T cell programs in steady state and inflammatory settings.**

A. Working model.

**Supplementary** **Table 1. Antibody information.**

| **Name** | **Catalog number** | **Company** | **Application** |
| --- | --- | --- | --- |
| RORγ pS203 / RORγt pS182 | 600-401-GR8 | Rockland-Inc | WB |
| BCL-xL | 2767S | Cell Signaling | Flow cytometry |
| RORγt | 12-6981-82 | eBiosciences | Flow cytometry |
| IL-10 | 12-7101-41 | eBiosciences | Flow cytometry |
| FOXP3 | 35-5773-82 | eBiosciences | Flow cytometry |
| CD3ε | 47-0033-82 | eBiosciences | Flow cytometry |
| CD11c | 15-1171-81 | eBiosciences | Flow cytometry |
| IL-17A | 506939 | BioLegend | Flow cytometry |
| IL-17F | 517006 | BioLegend | Flow cytometry |
| CD4 | 100410 | BioLegend | Flow cytometry |
| CD8α | 100723 | BioLegend | Flow cytometry |
| TCRγδ | 107503 | BioLegend | Flow cytometry |
| CD3e | 47-0033-82 | BioLegend | Flow cytometry |
| CD121a/IL1R | 113505 | BioLegend | Flow cytometry |
| F4/80 | 123109 | BioLegend | Flow cytometry |
| CD11b | 101226 | BioLegend | Flow cytometry |
| H3K27ac | 39133 | Active Motif | ChIP |
| RORγt | 14-6988-82 | LifeTech | ChIP |

**Supplementary Table 2. Primer sequences.**

| **qPCR primers** | **Sequences** |
| --- | --- |
| mCd4 F | CTTCGCAGTTTGATCGTTTTGAT |
| mCd4 R | CCGGACTGAAGGTCACTTTGA |
| mCd19_F | GGAGGCAATGTTGTGCTGC |
| mCd19_R | ACAATCACTAGCAAGATGCCC |
| mCd11b/Itgam_F | GCTCGACACCATCGCATCTA |
| mCd11b/Itgam_R | TGGTACTTCCTGTCTGCGTG |
| mCd11c/Itgax F | CTGGATAGCCTTTCTTCTGCTG |
| mCd11c/Itgax R | GCACACTGTGTCCGAACTCA |
| mCsf2 F | TCGTCTCTAACGAGTTCTCCTT |
| mCsf2 R | CGTAGACCCTGCTCGAATATCT |
| mGapdh F | AATGTGTCCGTCGTGGATCT |
| mGapdh R | CATCGAAGGTGGAAGAGTGG |
| mIfng F | ACAGCAAGGCGAAAAAGGATG |
| mIfng R | TGGTGGACCACTCGGATGA |
| mIl10 F | GCTGGACAACATACTGCTAACC |
| mIl10 R | ATTTCCGATAAGGCTTGGCAA |
| mIl12p40 F | TGCCAGGAGGATGTCACCT |
| mIl12p40 R | GGCGGGTCTGGTTTGATGAT |
| mIl17a F | TTTAACTCCCTTGGCGCAAAA |
| mIl17a R | CTTTCCCTCCGCATTGACAC |
| mIl17f F | TCCCCTGGAGGATAACACTG |
| mIl17f R | GGGGTCTCGAGTGATGTTGT |
| mIl1b F | GAAATGCCACCTTTTGACAGTG |
| mIl1b R | CTGGATGCTCTCATCAGGACA |
| mIl22 F | CCGAGGAGTCAGTGCTAAGG |
| mIl22 R | CATGTAGGGCTGGAACCTGT |
| mIl23a F | CAGCAGCTCTCTCGGAATCTC |
| mIl23a R | TGGATACGGGGCACATTATTTTT |
| mIl23r F | AGAGACACTGATTTGTGGGAAAG |
| mIl23r R | GTTCCAGGTGCATGTCATGTT |
| mIl6 F | TCTATACCACTTCACAAGTCGGA |
| mIl6 R | GAATTGCCATTGCACAACTCTTT |
| mTgfb F | CAACAATTCCTGGCGTTACC |
| mTgfb R | GCTGAATCGAAAGCCCTGTA |

| **ChIP-qPCR primers** | **Sequences** |
| --- | --- |
| mIl17a_P F | GGTGGTTCTGTGCTGACCTCATTT |
| mIl17a_P R | TGTGAGGTGGATGAAGAGTAGTGC |
| mIl17f_P F | CCTGGGTATGTCAAACAGCAGTAG |
| mIl17f_P R | GAGTAGAAACCCTGCATTCGTCAG |
| CNS2 F | TTGCATGCGCCTCCTAACACATAG |
| CNS2 R | TTTCTAGGTGGGTTCCTCACTGGT |
| CNS3a F | CAAGCATTCAGAAGGGAAGC |
| CNS3a R | CCATTCAGAAGGATCCCTGA |
| CNS3b F | CAGCTCACCACTGCCTGTAA |
| CNS3b R | GTCTGTGTGTGGGTTTGTGC |
| CNS4 F | TCGCCTCTTGACAAACAGTG |
| CNS4 R | TTCGTCCCTGTGATTTCCTC |
| CNS5 F | GTGAACGCAAGGGGTACAGT |
| CNS5 R | CCTTTGACCTTTCCCACAAA |
| Il23r_int8 F | TCCAGATTGCCTGACTGTATACCC |
| Il23r_int8 R | GAAGATGCACTTCTAGAAACCCGC |
